# Utilizing proteomics and phosphoproteomics to predict *ex vivo* drug sensitivity across genetically diverse AML patients

**DOI:** 10.1101/2021.06.04.447154

**Authors:** Sara JC Gosline, Cristina Tognon, Michael Nestor, Sunil Joshi, Rucha Modak, Alisa Damnernsawad, Jamie Moon, Joshua R. Hansen, Chelsea Hutchinson-Bunch, Marina A. Gritsenko, Karl K. Weitz, Elie Traer, Brian Drucker, Anupriya Agarwal, Paul Piehowski, Jason E. McDermott, Karin Rodland

## Abstract

Acute Myeloid Leukemia (AML) affects 20,000 patients in the US annually with a five-year survival rate of approximately 25%. One reason for the low survival rate is the high prevalence of clonal evolution that gives rise to heterogeneous sub-populations of leukemia. This genetic heterogeneity is difficult to treat using conventional therapies that are generally based on the detection of a single driving mutation. Thus, the use of molecular signatures, consisting of multiple functionally related transcripts or proteins, in making treatment decisions may overcome this hurdle and provide a more effective way to inform drug treatment protocols. Toward this end, the Beat AML research program prospectively collected genomic and transcriptomic data from over 1000 AML patients and carried out *ex vivo* drug sensitivity assays to identify signatures that could predict patient-specific drug responses. The Clinical Proteomic Tumor Analysis Consortium is in the process of extending this cohort to collect proteomic and phosphoproteomic measurements from a subset of these patient samples to evaluate the hypothesis that proteomic signatures can robustly predict drug response in AML patients. We sought to examine this hypothesis on a sub-cohort of 38 patient samples from Beat AML with proteomic and drug response data and evaluate our ability to identify proteomic signatures that predict drug response with high accuracy. For this initial analysis we built predictive models of patient drug responses across 26 drugs of interest using the proteomics and phosphproteomics data. We found that proteomics-derived signatures provide an accurate and robust signature of drug response in the AML *ex vivo* samples, as well as related cell lines, with better performance than those signatures derived from mutations or mRNA expression. Furthermore, we found that in specific drug-resistant cell lines, the proteins in our prognostic signatures represented dysregulated signaling pathways compared to parental cell lines, confirming the role of the proteins in the signatures in drug resistance. In conclusion, this pilot study demonstrates strong promise for proteomics-based patient stratification to predict drug sensitivity in AML.

## Introduction

Acute myeloid leukemia (AML) is characterized by the incomplete maturation of myeloblasts and their expansion in blood and bone marrow which impacts healthy blood cell formation resulting in decreased numbers of granulocytes, platelets, and red blood cells[1]. Even though the number of FDA-approved treatments for AML has increased significantly over the past five years, prognosis remains poor with a 5-year survival rate of 25% for individuals over the age of 20[2]. Targeted agents have shown promise in mutationally defined subsets of patients, but due to the genetic evolution of this highly heterogenous disease, drug response is often lost and patients relapse. Proper selection of personalized drugs and drug combinations over the course of a patient’s disease will be required to provide more durable clinical responses, and require a comprehensive mechanistic evaluation.

Computational modeling and machine learning approaches are able to predict the response of cancer samples to drug perturbation using baseline genomics or transcriptomics [3, 4]. This approach has been widely successful using data from the Cancer Cell Line Encyclopedia and subsequent dose response measurements carried out by the Broad and Sanger Institutes [5, 6] that identify specific signatures that predict which drugs affect cell lines from basal genomic and transcriptomic data of those same cell lines. These datasets have been further supplemented by global proteomic analysis of the same cell line library [7] that can also be used to predict drug response. However, cell line-derived computational models have their flaws, as they sample a limited subset of patient genetics and have been shown to correlate poorly with patient-derived xenograft data of the same tumor type [8]. There are still ongoing innovations in the computational space that predict drug response from underlying genomic phenotype [9] including Bayesian approaches [10], variational auto-encoders [11], and deep learning [12]. To date, however, most of these predictive models are based on cancer cell lines, which are limited in their ability to recapitulate the diversity of patient genetic backgrounds.

The Beat AML Dataset addresses the challenges of model systems by combining *ex vivo* small molecule inhibitor assays performed on freshly isolated patient leukemia cells with comprehensive genomic and transcriptomic data. In these studies, peripheral blood and bone marrow mononuclear cells (MNCs) from AML patients are isolated and exposed to a panel of approximately 145 drugs over a three-day period and cell viability is used as the primary readout for drug efficacy. Patient genomics and transcriptomics, as well as extensive clinical annotation, are also captured, enabling the stratification of patients by these measures [13]. The depth of sequencing performed on the samples allowed for the characterization of clonal architecture and enabled assessment of co-occurring mutational events, helping to identify drivers and co-actionable targets. This functional genomic dataset uncovered numerous novel genetic and microenvironmental drivers of AML pathogenesis and drug resistance, as well as corresponding validation of drug sensitivity profiles [14–18].

Through the National Cancer Institute’s Clinical Proteomic Tumor Analysis Consortium (CPTAC), patient-derived samples have been assayed using state-of-the-art mass spectrometry (MS) pipelines to produce proteomic and phosphoproteomic measurements of hundreds of tumors in breast, ovary, kidney, head and neck; endometrium, brain and other tissues [19–24]. In each study, these proteomic measurements reveal clinically relevant patterns that are not available at the genomic or transcriptomic level [25]; to date, however, these data have not been used to assess drug response in patient-derived tumors.

In this work, we combine the rigorous pre-clinical drug testing of the Beat AML dataset, together with patient-derived proteomic and phosphoproteomic measurements, to ascertain the potential for protein-level data to produce robust molecular biomarkers of drug response. We describe a retrospective study that uses the proteomic measurements of 38 patients to identify molecular signatures that can predict drug efficacy across a panel of 26 drugs of interest measured in our *ex vivo* protocol. We compare these signatures with those derived from genomic and transcriptomic data on the same patients and find proteomics to be extremely robust to inherent genetic sample diversity and often changes in abundance or phosphorylation of proteins that are directly related to the drug’s mechanism of action. We measured the ability of these signatures to predict drug response on numerous published hematopoietic cell lines and found them to provide high concordance (via rank correlation), if not perform better than, genomic or transcriptomic signatures. Lastly, we interrogated the expression of these signatures in two AML cell-line derived models of resistance to targeted therapeutic drugs, and identified dysregulated pathways that may play a potential role in mediating drug response.

## Experimental Procedures

### Experimental design and statistical rationale

Our overall experimental design is depicted in Figure S1. It entails the collection of patient AML samples, *ex-vivo* drug screening of these samples, and the construction of predictive models of drug response for each type of data collected.

### Sample collection

Samples were collected and processed as described in detail previously [13]. Briefly, all patients gave informed consent to participate in the Beat AML study, which had the approval and guidance of the Institutional Review Boards (IRB) from participating institutions. Mononuclear cells (MNCs) were isolated from fresh obtained bone marrow or peripheral blood samples from AML patients via Ficoll gradient centrifugation. Isolated MNCs were utilized for genomic (500x WES; RNA-seq) and *ex vivo* functional drug screens. WES and RNA-seq were performed using standard methods and data analysis was performed as previously described [13]. Clinical, prognostic, genetic, cytogenetic and pathologic laboratory values as well as treatment and outcome data were manually curated from the electronic medical records of the patient. Patients were assigned a specific diagnosis based on the prioritization of genetic and clinical factors as determined by WHO guidelines.

### *Ex vivo* drug screening analysis

For drug sensitivity assays, 10,000 viable cells were dispensed into each well of a 384-well plate containing 7 point series of drugs from a library of small molecule inhibitors. Cells were incubated with the drugs in RPMI media containing 10% FBS without supplementary cytokines. After 3 days of culture at 37 °C in 5% CO_2_, MTS reagent (CellTiter96 AQueous One; Promega) was added, the optical density was measured at 490 nm, and raw absorbance values were adjusted to a reference blank value and then used to determine cell viability (normalized to untreated control wells). *Ex vivo* functional drug screen data processing was performed as described [13], resulting quality controlled, probit fit drug curves which were used to calculate normalized AUC and IC50 values used in this analysis.

### Identifying drugs and samples for analysis

We selected 38 unique patients from our ongoing study that had complete proteomic and phosphoproteomic measurements. The list of available data for each patient is in **Table S1**. Although ~ 145 total compounds were tested in the drug panels, we filtered the drugs in this study to collect those that exhibited a range of responses across the 38 patients as determined by area under the curve (AUC) of the dose response. AUC represents the amount of drug required to reduce cell viability, so higher AUC mean the samples are less sensitive to the drug, and lower AUC indicates the samples are more resistance. Specifically, we selected drugs for which at least 10% or 2 (whichever was greater) samples exhibited an AUC less than 100 (determined to be sensitive to the drug in question); this definition produced a “balanced” distribution of AUC scores depicted in **Figure 1A**. We also added Gilteritinib (ASP-2215) to the panel as it is currently being evaluated in the clinic. The full range of drug responses across patients is shown in **Figure 1B**. While some patient samples lacked data on all 26 drugs (indicated in red in **Figure 1B**), we were still able to use these samples to compare the efficacy of genomics, transcriptomics, proteomics and phosphoproteomics to model drug sensitivity based on the available data.

**Figure 1:**
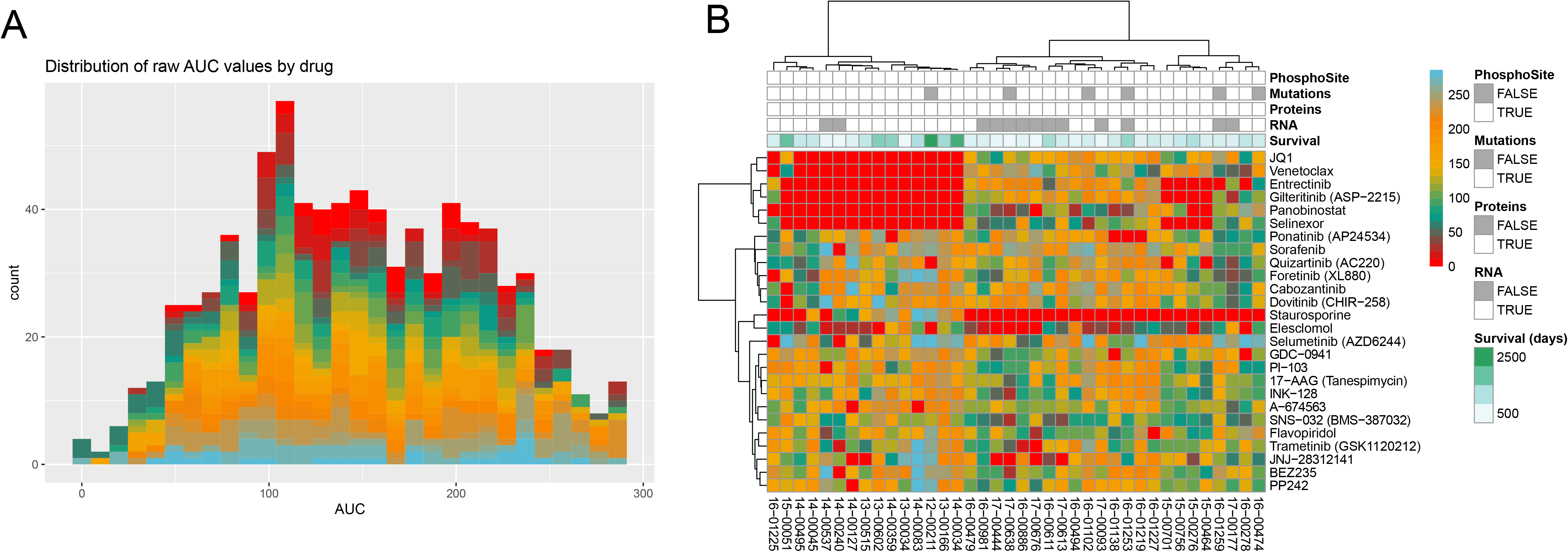
Summary of *ex-vivo* data collected using 38 primary AML samples. (A) Distribution of AUC values across the 26 drugs being evaluated. (B) Heatmap of AUC values across patient samples with the names of the drugs and the available data for each sample (grey bars, along top).

### Protein digestion and tandem mass tag (TMT) labeling

Sample preparation for proteomics was based on the protocol developed under the CPTAC consortium with minimal modifications[26]. Patient cell pellets were lysed with 500 μL fresh lysis buffer, containing 8 M urea (Sigma-Aldrich), 50 mM Tris pH 8.0, 75 mM sodium chloride, 1 mM ethylenediamine tetra-acetic acid, 2 μg/mL Aprotinin (Sigma-Aldrich), 10 μg/mL Leupeptin (Roche), 1 mM PMSF in EtOH, 10 mM sodium fluoride, 1% of phosphatase inhibitor cocktail 2 and 3 (Sigma-Aldrich), 20 μM PUGNAc, and 0.01 U/ μ/μL Benzonase. The samples were then vortexed for 10 seconds and then placed in thermomixer for 15 minutes at 4°C and 800 RPM, vortexing was then repeated and the samples incubated again for 15 minutes utilizing the same settings. After incubation, the samples were centrifuged for 10 minutes at 4°C and 18000 rcf to remove cell debris. The supernatant was then transferred to a fresh tube. A BCA (ThermoFisher) assay was performed on the supernatant to determine protein yield.

Protein concentrations were then normalized to the same concentrationprior to beginning digestion. The sample was reduced with 5 mM dithiothreitol (DTT) (Sigma-Aldrich) for 1 hour at 37°C and 800 rpm. Reduced cystines were alkylated with 10 mM iodacetamide (IAA) (Sigma-Aldrich) for 45 minutes at 25°C and 800 rpm in the dark. The sample was diluted fourfold with 50 mM Tris HCL pH 8.0 and then Lys-C (Wako) is added at a 1:20 enzyme:substrate ratio, followed by incubation for 2 hours at 25°C, shaking at 800 rpm. Trypsin (Promega) was then added at a 1:20 enzyme:substrate ratio, followed by a 14-hour incubation at 25°C and 800 rpm. The sample was quenched by adding formic acid to 1% by volume, and centrifuged for 15 minutes at 1500 rcf to remove any remaining cell debris. Peptides samples were desalted using a C18 solid phase extraction (SPE) cartridge (Waters Sep-Pak).

After drying down SPE eluates, each sample was reconstituted with 50 mM HEPES, pH 8.5 to a concentration of 5 μg/ μ/μL. Each isobaric tag aliquot was dissolved in 250 μL anhydrous acetonitrile to a final concentration of 20 μg/ μ/μL. The tag was added to the sample at a 1:1 peptide:label ratio and incubated for 1 hour at 25°C and 400 rpm and then diluted to 2.5 mg/mL with 50 mM HEPES pH 8.5, 20% acetonitrile (ACN). Finally, the reaction was quenched with 5% hydroxylamine and incubated for 15 minutes at 25°C and 400 rpm. The samples were then combined per each plex set and concentrated in a speed-vac before a final C18 SPE cleanup. Each 11-plex experiment was fractionated into 96 fractions by high pH reversed phase separation, followed by concatenation into 12 global fractions for MS analysis.

### LC-MS/MS analysis

Proteomic fractions were separated using a Waters nano-Aquity UPLC system (Waters) equipped with a homemade 75 um I.D. x 25 cm length C18 column packed with 1.9 um ReproSil-Pur 120 C18-AQ (Dr. Maisch GmbH). A 120-minute gradient of 95% mobile phase A (0.1% (v/v) formic acid in water) to 19% mobile phase B (0.1% (v/v) FA in acetonitrile) was applied to each fraction. The separation was coupled to either a Thermo Orbitrap™ Fusion Lumos™ (patient samples) or Q Exactive™ HF (cell lines) Hybrid Quadrupole-Orbitrap™ mass spectrometer for MS/MS analysis. MS Spectra were collected from 350 to 1800 m/z at a mass resolution setting of 60,000. A top speed method was used for the collection of MS2 spectra at a mass resolution of 50K. An isolation window of 0.7 m/z was used for higher energy collision dissociation (HCD), singly charged species were excluded, and the dynamic exclusion window was 45 seconds. For the Fusion Lumos™, a top speed method was used for the collection of MS2 spectra at a mass resolution of 50K. For the Q Exactive™,™ HF, experiments a top 16 method was used for the collection of MS^2^ spectra at a mass resolution of 30K.

### TMT global proteomics data processing

All Thermo RAW files were processed using mzRefinery to correct for mass calibration errors, and then spectra were searched with MS-GF+ v9881[27–29] to match against the human reference protein sequence database downloaded in April of 2018 (71,599 proteins), combined with common contaminants (e.g., trypsin, keratin). A partially tryptic search was used with a ± 10 parts per million (ppm) parent ion mass tolerance. A reversed sequence decoy database approach was used for false discovery rate calculation. MS-GF+ considered static carbamidomethylation (+57.0215 Da) on Cys residues and TMT modification (+229.1629 Da) on the peptide N terminus and Lys residues, and dynamic oxidation (+15.9949 Da) on Met residues. The resulting peptide identifications were filtered to a 1% false discovery rate at the unique peptide level. A sequence coverage minimum of 6 per 1000 amino acids was used to maintain a 1% FDR at the protein level after assembly by parsimonious inference.

The intensities of TMT 11 reporter ions were extracted using MASIC software[30]. Extracted intensities were then linked to PSMs passing the confidence thresholds described above. Relative protein abundance was calculated as the ratio of sample abundance to reference channel abundance, using the summed reporter ion intensities from peptides that could be uniquely mapped to a gene. The relative abundances were log2 transformed and zero-centered for each gene to obtain final relative abundance values.

### TMT phosphoproteomics data processing

IMAC enriched fraction datasets were searched as described above with the addition of a dynamic phosphorylation (+79.9663 Da) modification on Ser, Thr, or Tyr residues. The phosphoproteomic data were further processed with the Ascore algorithm[31] for phosphorylation site localization, and the top-scoring assignments were reported. To account for sample loading biases in the phosphoproteome analysis, we applied the same correction factors derived from median-centering of the global proteomic dataset for normalization.

All proteomic data can be found on our synapse site. The cohort is spread across three tranches, as described below.

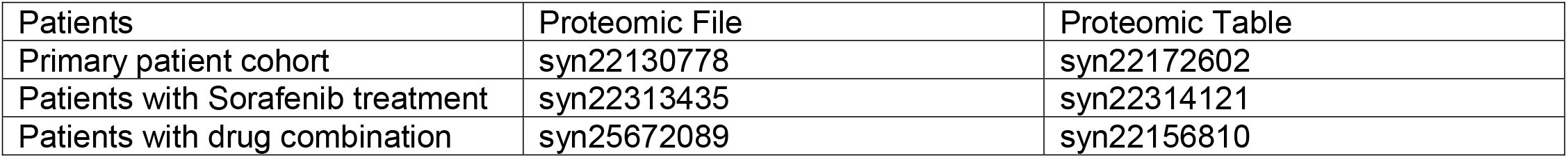

| Patients | Proteomic File | Proteomic Table |
| --- | --- | --- |
| Primary patient cohort | syn22130778 | syn22172602 |
| Patients with Sorafenib treatment | syn22313435 | syn22314121 |
| Patients with drug combination | syn25672089 | syn22156810 |

The phophoproteomic measurements are also divided, with the resources listed below.

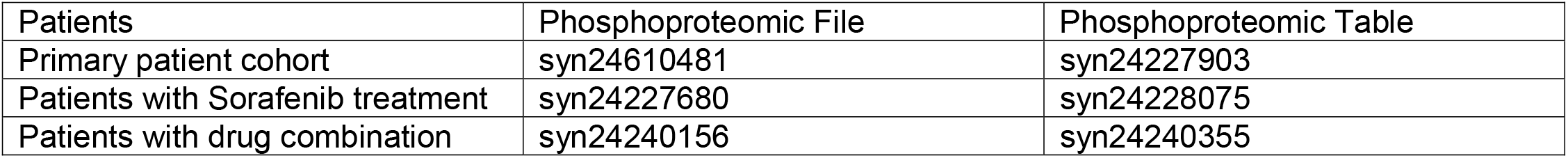

The phosphoproteomic measurements are also divided, with the resources listed below.
| Patients | Phosphoproteomic File | Phosphoproteomic Table |
| --- | --- | --- |
| Primary patient cohort | syn24610481 | syn24227903 |
| Patients with Sorafenib treatment | syn24227680 | syn24228075 |
| Patients with drug combination | syn24240156 | syn24240355 |

### Linear models of proteomics and drug response

We constructed linear models for each of the 26 different drugs across up to 38 patients (depending on how many patient samples were evaluated with that drug) by regressing the AUC values (which ranged between 0 and 300, as depicted in **Figure 1A)** against the molecular data as shown in **Table S1**. The input data for each model were each scaled slightly differently: the genetic mutations were represented as a binary matrix in which a 1 represented the presence of a somatic mutation and a 0 represented no mutation, the transcriptomics was represented by Counts per million (CPM) of gene expression values, while proteomics and phosphoproteomics were represented as the log ratio of gene/phosphosites compared to the reference sample described above.

For each data type/drug combination, we constructed a linear model **Y**~**X** where **Y** represents the vector of AUC values and **X** represents the molecular measurements for that patient. To reduce the number of features selected by the model we used the LASSO regression [32] as implemented by the ‘glmnet’ package [33]. We employed leave one out cross-validation for each combination of data to select the alpha parameter that minimized cross-validation error. All of our analysis can be found in the ‘amlresistancenetworks’ package we built at http://github.com/PNNL-CompBio/amlresistancenetworks and implemented at https://github.com/PNNL-CompBio/beatamlpilotproteomics. Those models that failed to select any molecular features were not included in our final analysis, depicted in **Table S2**. To evaluate the hierarchical clustering of the features selected by the models, we used the ‘pheatmap’ R package based on the log ratio values for the proteomics and phosphoproteomics features selected. The results are depicted in **Figure 2** and summarized in **Table 1**.

**Figure 2:**
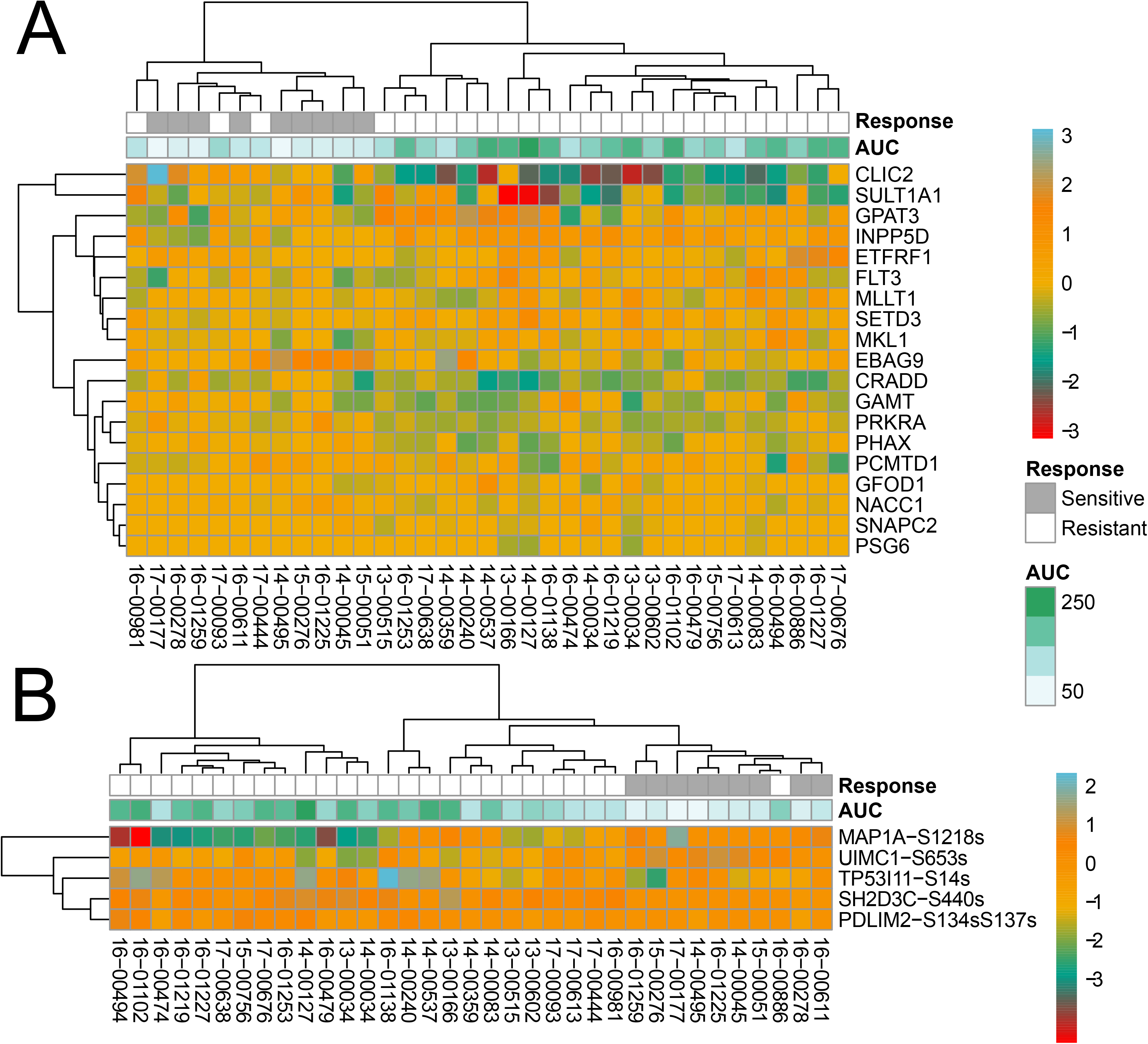
LASSO model of quizartinib response. (A) Expression of proteins selected by the LASSO model of proteomics measurements. (B) Expression of phosphosites selected by the LASSO model of phosphosite measurements.

**Table 1:** Summary of LASSO classifier performance. For each data type, we describe the total number of drugs modeled, the mean standard error, the mean correlation (R) and the mean number of samples used to build each model.

| DATA TYPE | MODELS | MEAN<br>SQUARED<br>ERROR | R | SAMPLES |
| --- | --- | --- | --- | --- |
| BINARY<br>MUTATIONS | 21 | 2889.8 | 0.84 | 27.38 |
| MRNA LEVELS | 12 | 3659.7 | 0.95 | 19.17 |
| PHOSPHOSITE | 21 | 3198.8 | 0.88 | 33.05 |
| PROTEIN LEVELS | 20 | 2728.7 | 0.90 | 33.3 |

We also discreted the AUC by representing **Y** as a binary variable, where 1 represented an AUC greater than 100 (patient is resistant to drug) and 0 if the AUC is less than 100 (patient is sensitive to drug) and used this as input into a logistic regression using ‘glmnet’. Example results depicted in **Figure 3** and summarized in **Table 2**. Direct comparisons of the LASSO and Logistic regression models are shown in **Figure 4**.

**Figure 3:**
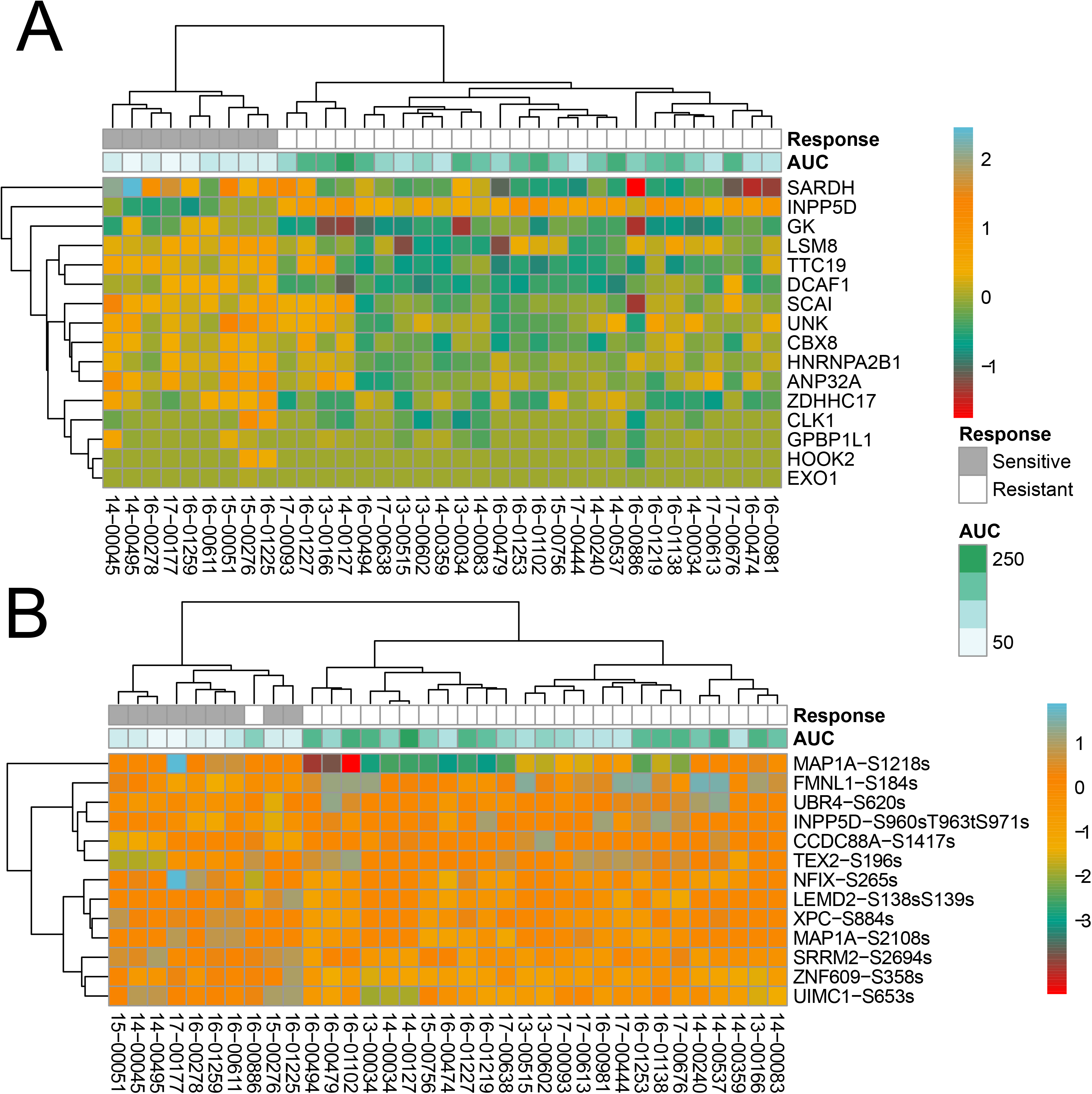
Logistic model of quizartinib response assessment cross-validation performance across data modalities. (A) Expression of proteins selected by the logistic model. (B) Expression of phosphosites selected by the logistic model.

**Table 2:** Summary of Logistic classifier performance. For each data type, we describe the total number of drugs modeled, the mean misclassification error, the mean correlation (R) and the mean number of samples used to build each model.

| MEASUREMENT | NUMBER MODELS | MISCLASSIFICATION ERROR | MEAN R | SAMPLES |
| --- | --- | --- | --- | --- |
| **BINARY MUTATIONS** | 13 | 0.29 | 0.84 | 28.15 |
| **MRNA LEVELS** | 13 | 0.34 | 0.74 | 20.69 |
| **PHOSPHOSITE** | 15 | 0.33 | 0.73 | 33.73 |
| **PROTEIN LEVELS** | 17 | 0.28 | 0.69 | 33.94 |

**Figure 4:**
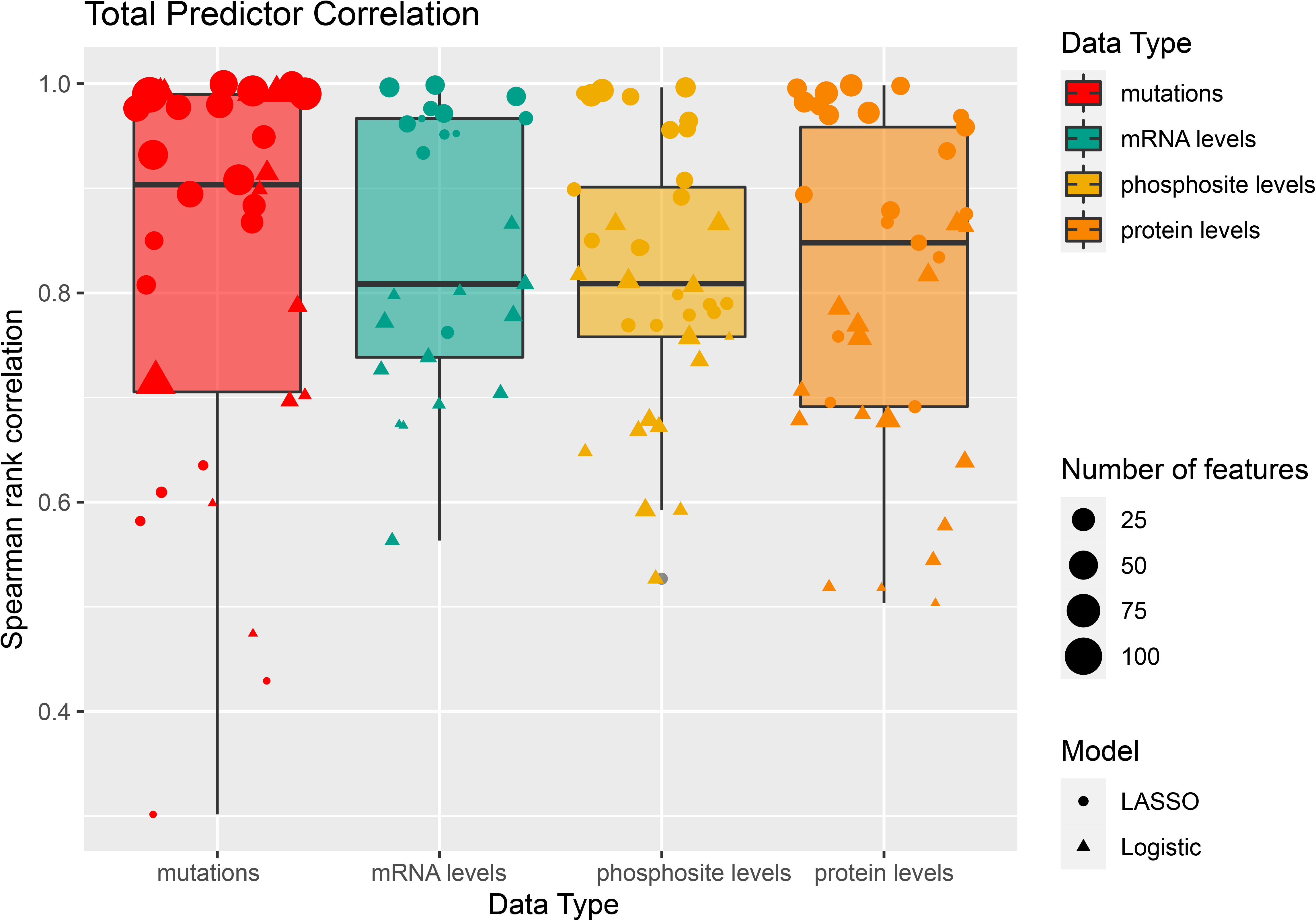
Comparison of LASSO (circle) and Logistic (triangle) models across data types, measured by Spearman rank correlation.

### Signature interpretation using pathway annotations and statistical enrichment

To identify patterns in the features selected by the LASSO and Logistic models we employed three main approaches. For gene, transcript, and proteomic signatures, we first used the ‘clusterProfiler’ package [34] to identify GO biological process tools that are enriched for the specific genes, transcripts, or proteins selected by the model. The results are listed in **Table S2**. In cases where there were no significant (corrected p<0.01) terms, the column is blank. For phosphoproteomic features, we used the ‘leapR’ R package[35] to identify specific kinases that were over-represented among the selected substrates, though none were identified. We believed this is due to the sparsity of the signatures as well as the lack of more information about the kinase-substrate interactions.

### Cell line data comparison

To evaluate the patient-derived signatures on an external dataset we collected mRNA expression data from and mutational data from the Cancer Cell Line Encyclopedia (CCLE)[36] together with proteomics data from the same cell lines [7]. The data is merged into a single Synapse table at https://www.synapse.org/#!Synapse:syn23674921/tables/. Drug sensitivity data was collected from both the Cancer Therapeutics Response Portal (CTRP) [5, 37] and Sanger [6] datasets and stored at https://www.synapse.org/#!Synapse:syn23004543/tables/.

Cell lines were filtered for those from hematopoietic origin, to include the 22 showed in **Table S3**. Of the 25 drugs we modeled using the Beat AML patient cohort, we identified 17 which were also measured in cancer cell lines. We therefore took the LASSO and Logistic models we built for these 17 drugs and evaluated their performance on the cell line data. We applied a linear transformation to the AUC data from each dataset to align with the AUC values from our patient cohort. The performance of each predictive model is depicted in **Table S4** and **Figure 5**.

**Figure 5:**
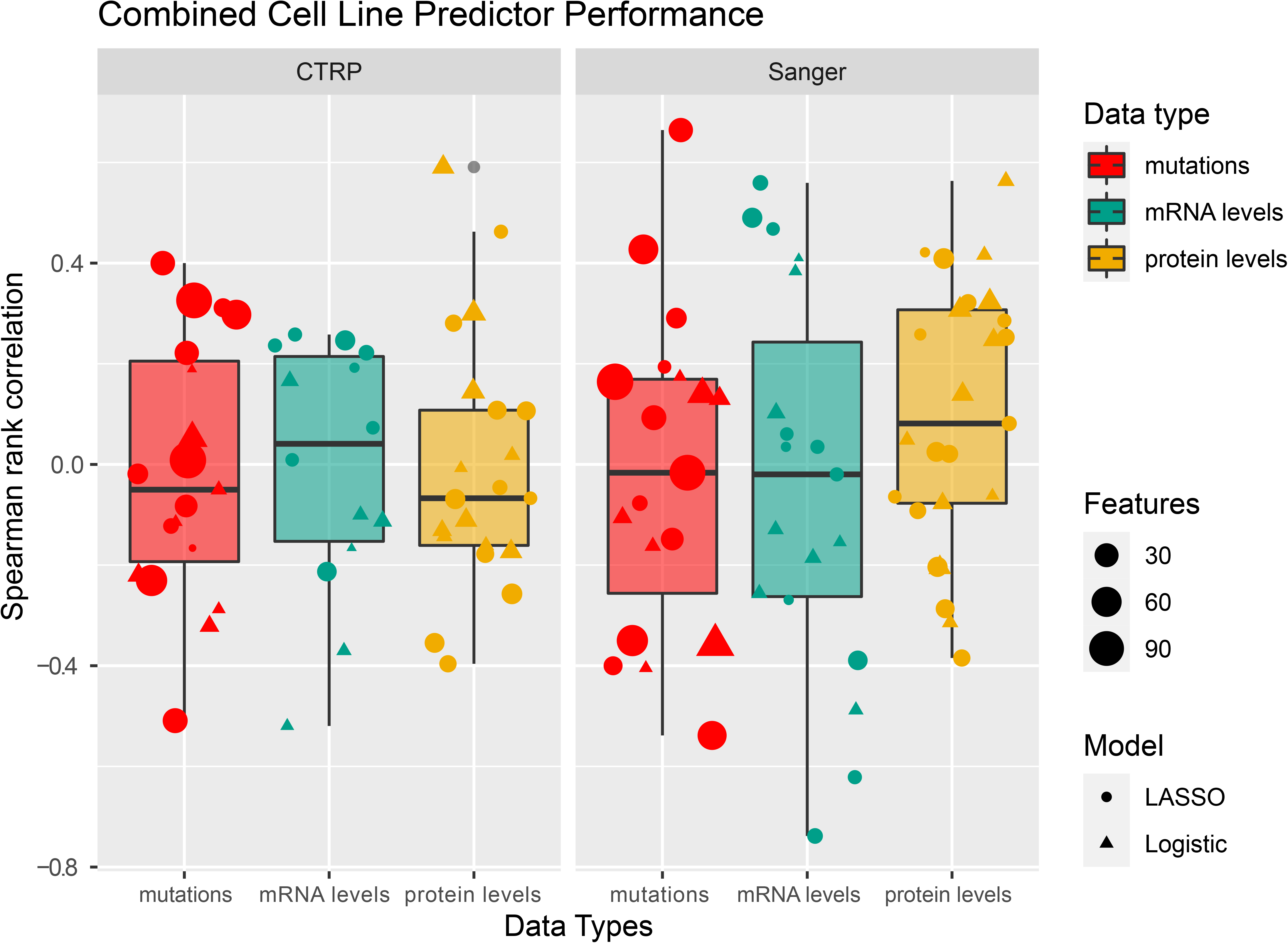
Comparison of LASSO (circle) and Logistic (triangle) models across data types, measured by Spearman rank correlation.

### Trametinib resistant cell line cultures

Human MOLM-13 cells with FLT-ITD mutation, were obtained from the Sanger Institute Cancer Cell Line Panel. Cell lines were maintained in RPMI 1640 (Gibco) supplemented with 20% Fetal Bovine Serum (HyClone), 2% L-glutamine, 1% penicillin/streptomycin (Life Technologies). Trametinib-resistant MOLM13 cell lines were generated by culturing MOLM13 cells in increasing concentrations of trametinib (Selleck). Cell viability was measured bi-weekly and cells were replenished with new media and trametinib. Resistance was assessed using the MTS assay for drug sensitivity. Once resistant, cell lines were maintained in 50nM trametinib added bi-weekly. Cell lines were screened for mycoplasma contamination on a monthly schedule.

For proteomic and phosphorproteomic profiling, 5 million parental MOLM13 (N=3) and resistant MOLM13 cell lines (N=3) cell lines were starved overnight in starvation media (RPMI supplemented with 0.1% BSA). Trametinib (50nM) was added to the starvation media of the resistant cell lines. Cells were washed thrice in PBS, pelleted and flash frozen.

### Quizartinib resistant cell line cultures

Human MOLM14 cells were generously provided by Dr. Yoshinobu Matsuo (Fujisaki Cell Center, Hayashibara Biochemical Labs, Okayama, Japan). Cells were grown in RPMI (Life Technologies Inc., Carlsbad, CA) supplemented with 10% FBS (Atlanta Biologicals, Flowery Branch, GA), 2% L-glutamine, 1% penicillin/streptomycin (Life Technologies Inc.), and 0.1% amphotericin B (HyClone, South Logan, UT). Cell line authentication was performed at the OHSU DNA Services Core facility.

To establish resistant cultures, 10 million MOLM14 cells were treated with 10 nM of quizartinib (Selleck Chemicals, Houston, TX) in media alone (N = 4) or in media supplemented with 10 ng/mL of FGF2 (N = 4) or FLT3 ligand (N = 4, FL; PeproTech Inc., Rocky Hill, NJ)[38]. All cultures were maintained in 10 mL of media. Every 2 or 3 days, recombinant ligands and quizartinib were replaced and cell viability was evaluated using the Guava personal flow cytometer (Millipore Inc., Burlington, MA). Following ligand withdrawal, quizartinib and media were similarly replenished and viability was monitored every 2 to 3 days. All cell lines were tested for mycoplasma on a monthly schedule.

For proteomic and phosphoproteomic profiling, naïve MOLM14 (N = 4), quizartinib-resistant parental (N = 2, no ligand), early (N = 4/ligand) and late (N = 4/ligand) cultures were washed three times with PBS to remove any trace of fetal bovine serum, pelleted, and flash frozen.

### Building networks from proteomic signatures and network reduction strategies

To provide further context for the phosphoproteomic features selected by the models, we mapped selected phosphosites and proteins to published protein-protein [39] and kinase-substrate[40, 41] interactions and then reduced this network to identify subnetworks using the ‘PCSF’ R package [42, 43]. Specifically, we used the STRING database [39] together with networkKin[40] and PhosphoSitePlus[41] predictions of kinase substrate interactions to build a graph that combined protein-protein interactions with kinase-substrate interactions. To do this we added each phosphosite as its own node in the underlying graph. We weighted each edge from the node representing the substrate gene to the phosphosite with a cost of *m*/4 where *m* represents the mean cost of all the edges in the graph. The weight of each edge between the phosphosite node and the kinase gene was weighted with a cost of 3*m*/2 where *m* represents the mean cost of all edges in the graph. We then ran the PCSF algorithm[42, 43] over 100 randomizations using phosphosites together with proteins from a single drug model. The results for the Trametinib LASSO signatures are in **Figure 6C**, and the results for the Quizartinib LASSO signatures are in **Figure 7C**.

**Figure 6:**
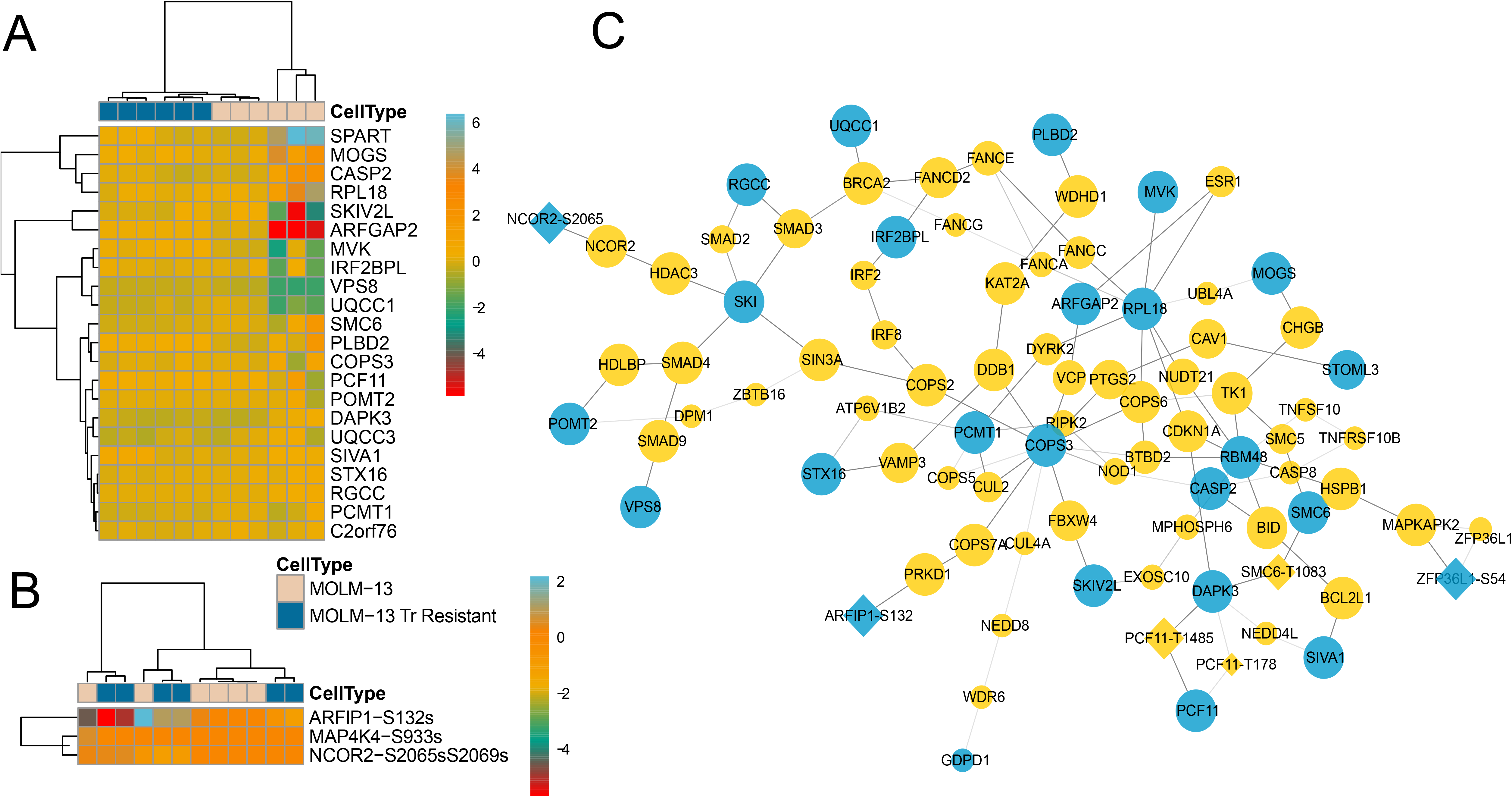
Expression and role of LASSO-derived proteins and phosphosites in trametinib resistant cell lines. (A) Clustering of proteins selected by the regression model in parental MOLM-13 (beige squares) and trametinib-resistant MOLM-13 (blue) cells. (B) Clustering of phosphosites selected by the LASSO model in the same cell lines. (C) Proteins (circles) and phosphosites (diamonds) selected by the prize-collecting Steiner forest algorithm based on the presence of the blue proteins/substrates in the protein signature. Yellow nodes of the graph are selected to form a single cohesive network

**Figure 7:**
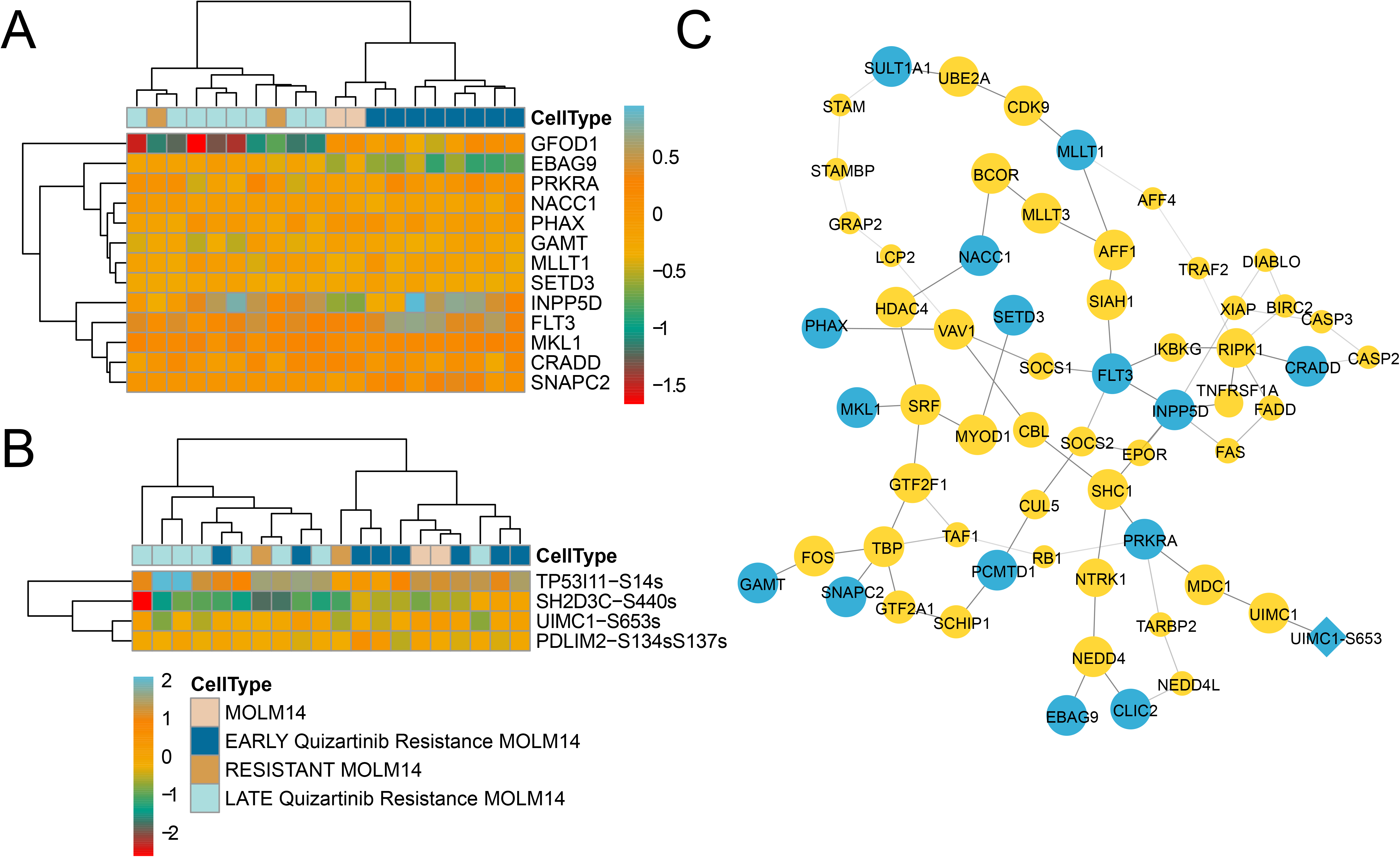
Expression and role of LASSO-derived proteins and phosphosites in Quizartinib resistant cell lines. (A) Clustering of proteins selected by the regression model in parental MOLM-14 (beige squares) and quizartinib-resistant MOLM-14 (blue) cells. (B) Clustering of phosphosites selected by the LASSO model in the same cell lines. (C) Proteins (circles) and phosphosites (diamonds) selected by the prize-collecting steiner forest algorithm based on the presence of the blue proteins/substrates in the protein signature. Yellow nodes of the graph are selected to form a single cohesive network

Using the proteins selected by the PCSF algorithm, which are a combination of those selected by the linear model as well as those selected by the PCSF algorithm, we used Cytoscape [44] and the BinGO [45] application to identify which GO biological process terms were enriched. The results are depicted in **Tables S5** and **S6**.

## Results

### Linear models identify robust proteomic signatures of drug response

We constructed linear models as described above for each combination of drug and data modality for total of 104 different combinations. Due to data loss (e.g., lack of drug response data) and general lack of signal, we were able to build LASSO models for 74 of these combinations, including 20 and 21 models using proteomic and phosphoproteomic data, respectively. To assess the quality of these models we employed both quantitative and qualitative metrics.

Our quantitative signature metrics are summarized in the bottom two rows of **Table 1**. The ideal data modality would be able to model the most drugs (second column) with the lowest amount of mean squared error (third column) and the highest correlation with the actual value (R, fourth column). The proteomic and phosphoproteomic data can model most of the drugs with low mean error and high mean correlation.

To assess the signatures in a qualitative manner, we used two other metrics. First, we used hierarchical clustering to determine if the values of the model-selected features clustered the patient samples in a way that recapitulated drug response. An example of this approach on quizartinib is shown in **Figure 2B and C**. Here we were able to identify proteins (**Figure 2A**) and phosphosites (**Figure 2B**) whose expression clustered patient samples by the AUC of Quizartinib in the *ex vivo* patient samples using the LASSO regression. Specifically, we demonstrate that the expression of FLT3, a direct target of quizartinib, is reduced in samples that are more sensitive to quizartinib (**Figure 2A**).

As a second qualitative measure, we tested the features selected by the model to ask if specific GO biological processes or kinases were enriched in the proteins (**Figure 2A**) or phosphosites (**Figure 2B**) selected by the models. Of the 20 LASSO proteomic regression models, 14 had at least one significantly enriched biological process. The results for these signatures (and all others) are depicted in **Table S2**.

### Logistic regression predicts fewer drug responses with less accuracy

Given the sparsity of the data we sought to explore the ability to create a binary predictor of drug response using a logistic regression model [33]. Across the 104 drug/data combinations, we were able to build logistic regression models for 57, fewer than what was possible for the LASSO. Of those, only 17 and 15 drugs using proteomic and phosphoproteomic data.

We evaluated the logistic proteomic regression models using the same quantitative and qualitative metrics as for the linear regression. The summary statistics are depicted in **Table 2** and the hierarchical clustering in **Figure 3**. The summary statistics are similar to those in **Table 1**, with the exception of the error metric, as classification models are measured via a misclassification error (fraction of samples misclassified) instead of mean squared error. **Figure 3** illustrates the proteomic and phosphosite features selected to predict the quizartinib response, which exhibited tight clusters using proteins selected by the logistic model. Again these heatmaps enabled us to further interrogate the proteins identified. In Figure 3 we noticed INPP5D, which is identified in both the LASSO and logistic regression models and highly down-regulated in sensitive samples (**Figure 3A**). This gene encodes the inositol 5-phophatase know as SHIP1 which acts as a negative regulator of the PI3K/AKT pathway. SHIP-1 affects cell proliferation in AML, due to mutational alteration of the nuclear localization signature or phosphorylation [46]. It has also been shown to act as an adaptor protein linking to wild type FLT3 signaling[47, 48]. Lastly, we also evaluated GO enrichment, and found that fewer (7 of the 17) proteomic signatures selected by the logistic regression had at least one enriched biological process when compared to the LASSO.

We compared the overall performance of proteomic and phosphoproteomic derived models (described above) to those derived from mutational or transcriptomic profiles. Many drugs, such as quizartinib or trametinib, a Mek kinase inhibitor, are designed to target important signaling pathways found to be frequently hyperactivated by specific mutations and therefore it is expected that the presence/absence of mutations in the FLT3/ MAPK pathway (FLT3-ITD, FLT3-TKD, RAS, PTPN11, NF1) will impact the efficacy of the drug [13]. Furthermore, transcriptional signatures derived from regression models have also proven to be able to predict drug response [13]. As such, we wanted to determine if proteomic features, which are more removed from direct genetic mutations but more proximal to function, are as effective in predicting drug response.

We compared the two regression approaches to each other using the correlation metric, depicted in **Figure 4**. In short, the LASSO regression is superior to the Logistic in terms of overall correlation across all four data modalities and was also able to model more drugs, as indicated by the number of dots compared to triangles. When comparing data types used as input to the model, the proteomic and phosphoproteomic models performed as well as or better than those derived from mRNA or genomic data. While genomic mutation-derived models were generally more correlated with the training dataset (**Figure 4**), they were far less accurate in terms of mean squared error (**Figure S2A)** and misclassification error (**Figure S2B**).

Since this patient cohort was primarily focused on the assessment of proteomic and phosphoproteomic measurements, we recognize that our results could be biased toward protein-level data due to larger training sets (**Table S1**, **Figure 1B**). We therefore repeated our analysis for a “square” dataset using the 18 samples that had all four data types and built our models of drug response on these patients. While highly limited, these results, depicted in **Figure S3**, show that proteomics and phosphoproteomics were able to produce performant signatures using leave-one-out cross validation, with higher correlation values in models built with proteomic and phospho-proteomic data than those from genomic or transcriptomic data.

### Comparison with hematopoietic cell line data suggests proteomic signatures are most robust

Given the limited data available for cross validation in the Beat AML cohort as well as the lack of ability to control for genetic heterogeneity, we sought existing cell line panels as an external validation set for our prognostic signatures. We used the signatures developed in the Beat AML patient cohort and evaluated their ability to predict drug response in two publicly available cell line drug sensitivity studies (measuring percent viable cells post drug treatment), using genetic mutations, gene expression values, and proteomic data. Specifically we collected hematopoetic cell line data from the Cancer therapeutics response portal (CTRP) [5] and Sanger [6] studies and measured the predictions on 17 drugs for which we had valid patient signatures and also data in cell lines from at least one of the two studies, described in **Table S5**. The results, depicted in **Figure 5**, show that, while only a handful of signatures were successfully applied to the cell line data, they generally correlated at least as well if not better than models built from genomic or transcriptomic data, and also had lower error (**Figure S3**).

### Proteins and phosphosites that predict trametinib response are dysregulated in resistant cell lines

Given that the proteins and phosphosites selected by our models were able to distinguish patient samples that responded to drugs from those that did not, we hypothesized that the proteins selected by the models play a distinct role in drug response. To test this, we sought to alter the drug response *in vitro* by culturing AML cell lines in the presence of low concentration of a drug we modeled and measuring how the proteome changed in resistance samples. We focused our study one trametinib, a drug that targets the MAPK signaling cascade, over 3-4 months and compared the expression of the model-selected proteins in response to trametinib treatment.

We clustered the expression values of the proteins selected by the patient-derived LASSO trametinib regression model in both the trametinib-resistant and parental cell lines (across multiple replicates) and found two distinct clusters, depicted in **Figure 6A,** that aligned with the drug-resistant phenotype. When we clustered the phosphosites selected by the same LASSO model, it also grouped many parental cell lines in the same cluster (**Figure 6B**). While many drug signatures did not exhibit significant GO enrichment (**Table S2**), we found two biological processes to be enriched with proteins from the trametinib LASSO signature: respiratory chain complex III assembly and mitochondrial respiratory chain complex III assembly. The limited number of enriched GO terms is driven by the small size of the LASSO regression model.

Given the limited number of GO terms identified by the proteins selected in this model, we expanded the proteins and phosphosites selected by the model by building a network model that joined the selected proteins and phosphosites in a network using published protein-protein [39] and kinase-substrate[40, 41] interactions (see Methods) via the Prize-Collecting Steiner Forest algorithm [42, 43]. The result, depicted in **Figure 6C**, highlights the proteins from our statistical model in blue together with neighboring proteins in yellow. Using this enhanced network model we were able to identify more significantly over-represented GO terms than by the model-selected proteins along. This analysis, depicted in **Table S5**, shows that many apoptosis and cell death-related pathways are over-represented, primarily driven by the presence of known apoptotic genes DAPK3 and CASP2 in the proteomics signature in **Figure 6A**.

### Proteins and phosphosites that predict quizartinib response are dysregulated in models of late, but not early, resistance

To further explore the extent to which the protein signatures affect drug response, we utilized a recently-published cell line model that represents patients that develop early vs late resistance to the FLT3 inhibitor quizartinib. Here, cells are cultured in the presence of a quizartinib plus an activating ligand – either the FLT3 ligand (FL) or FGF2[49]. Using this model we can compare cell lines that show symptoms of early resistance (shortly after co-treatment) or late resistance, which is ligand independent and accompanied by the outgrowth of genetically altered resistant clones[49]. We hypothesized that the patient-derived signature would more closely resemble the long-term resistance phenotype.

To test this hypothesis, we plotted the proteins and phosphosites selected by the LASSO model in these cell lines, as depicted in **Figures 7A** and **7B** respectively. We observed a similar split between sensitive and resistant cells as we did in Figure 6, as the proteins and phosphosites that predict drug response cluster the MOLM14 parental cells (beige) distinctly from the fully resistant cells (gold). However, in this case, these proteins separates those cells that represented ‘early resistance’ (dark blue) from those that represent ‘late resistance’ (light blue) in our previous work. This fits with our previous claim that the resistance to FLT3 inhibitors involves a two-step process, as cell lines exhibiting the early resistance phenotype cluster more closely to the parental cells than to the late resistance cells. Since no GO terms appeared to be enriched among the proteins in the LASSO signature (**Table S2**), we also examined the network linked by the proteins and phosphosites using the Prize-Collecting Steiner Forest [42, 43] as described in our Experimental Procedures and depicted in **Figure 7C**. Because the proteins and phosphosites selected by the model were not enriched in any specific GO terms, the network enabled us to get a broader picture of how the proteins involved could participate in the same signaling pathways. Similar to the trametinib network, these proteins were also enriched in apoptotic related pathways (**Table S6**), but unlike the previous network we found enrichment in B cell activation, differentiation, and homeostasis, driven by network proteins INPP5D, FLT3, and CASP3.

## Discussion

This study describes a new approach to predicting drug response in AML patient samples using global proteomic and phosphoproteomic data in combination with traditional genomic and transcriptomic approaches. Both linear regression (LASSO) and discrete (binarized) logistic regression approaches were used on a set of samples from 38 AML patients with corresponding *ex vivo* drug response data. Both LASSO and logistic regression approaches performed satisfactorily, with quantitative metrics of model coverage, error and correlation comparable between mutation-based, transcript-based, protein-based and phosphosite-based approaches. Neither LASSO nor logistic regression was clearly superior, with LASSO providing slightly better quantitative metrics while the logistic regression model was more robust when applied to cell line data. These positive results suggest that protein biomarkers could be used to better stratify patients to identify which treatment would be best for their disease.

Models generated on patient samples were tested for performance on both published cell line perturbation datasets (CTRP and Sanger), and additionally on two in-house models of drug resistance based on AML cell lines under prolonged inhibitor exposure. The predictive ability of the proteomic and phosphosite models was quite strong, generating tight clusters of sensitive and resistant cell lines, and effectively subdividing cell lines treated with stromal factors to model early and late drug resistance into their respective phases. Use of Prize-Collecting Steiner Forest algorithms to identify specific protein networks that were altered to gene set enrichment based on GO ontologies alone, highlighting apoptotic pathways that have been shown to be synergistic with both trametinib and quizartinib. Recent drug trials of venetoclax, a drug that targets the apoptotic pathway, has been shown to be successful in combination with trametinib[50] and quizartinib[51].

In summary, this study presents an effective workflow for the future analysis of integrated genomic, transcriptomic, proteomic and phosphoproteomic data in larger cohorts, such as the 210 patient Beat AML cohort currently under analysis. While the patient cohort used in this preliminary study is relatively limited in size, the robust verification in cell line studies provides confidence in the scalability of the method. Additionally, the comparable performance of protein-based models compared to mutation-based models opens up the possibility of developing antibody-based, CLIA-eligible assays for the rapid assessment of likely therapeutic targets at the time of biopsy or surgery, without the need for DNA sequencing. Lastly we believe that our network approaches could identify other potential drug synergies that have not yet been tried in the clinic.

We believe that studying the proteins and phosphosites directly can enable new biological insights into the mechanisms of drug response and resistance. This has been shown in both *in vitro* studies and studies with archived patient samples (i.e., CPTAC studies), as measuring proteins directly can identify the dysregulation in signaling beyond the genetic mutations or altered transcripts.

## Abbreviations

AML: Acute Myeloid Leukemia
CPTAC: Clinical Proteomic Tumor Analysis Consortium
FDR: False Discovery Rate
AUC: Area Under the Curve

## Data Availability

Data was uploaded to Synapse where it was used for subsequent analysis at http://synapse.org/ptrc. mRNA (counts per million) and genetic mutation measurements (variant allele frequency) can be found at https://www.synapse.org/#!Synapse:syn22172602/tables/.

All data used for this project is stored on Synapse at http://synapse.org/ptrc, where you can request access to the data specifically mentioned in this manuscript. All analysis and figures can be viewed at https://github.com/PNNL-CompBio/beatamlpilotproteomics.

## Supplemental Tables

**Supplemental Table 1**: List of drugs and available samples

**Supplemental Table 2**: All identified signatures and their functional enrichment (GSEA/KSEA)

**Supplemental Table 3**: Description of cell lines used

**Supplemental Table 4**: Number of cell line measurements on our drug panel

**Supplemental Table 5**: GO enrichment of Trametinib network

**Supplemental Table 6**: GO enrichment of Quizartinib network

## Supplemental Figures

**Supplemental Figure 1**: Overview figure describes the experimental design.

**Supplemental Figure 2:** Error rate calculations across predictors within patient samples.(A) Mean squared error across LASSO models, (B) Misclassification error across Logistic models

**Supplemental Figure 3:** Error rate calculations across predictors with equal numbers of samples for each data type.(B) Mean squared error across LASSO models, (C) Misclassification error across Logistic models

**Supplemental Figure 4**: Error rate calculations across predictors on cell line data..(A) Mean squared error across LASSO models, (B) Misclassification error across Logistic models

## Notes

https://github.com/PNNL-CompBio/beatAMLpilotProteomics

http://synapse.org/ptrc

### Competing Interest Statement

The authors have declared no competing interest.

