## Supplementary figures and images for "Utilizing proteomics and phosphoproteomics to predict *ex vivo* drug sensitivity across genetically diverse AML patients"

### Supplemental Figure 1

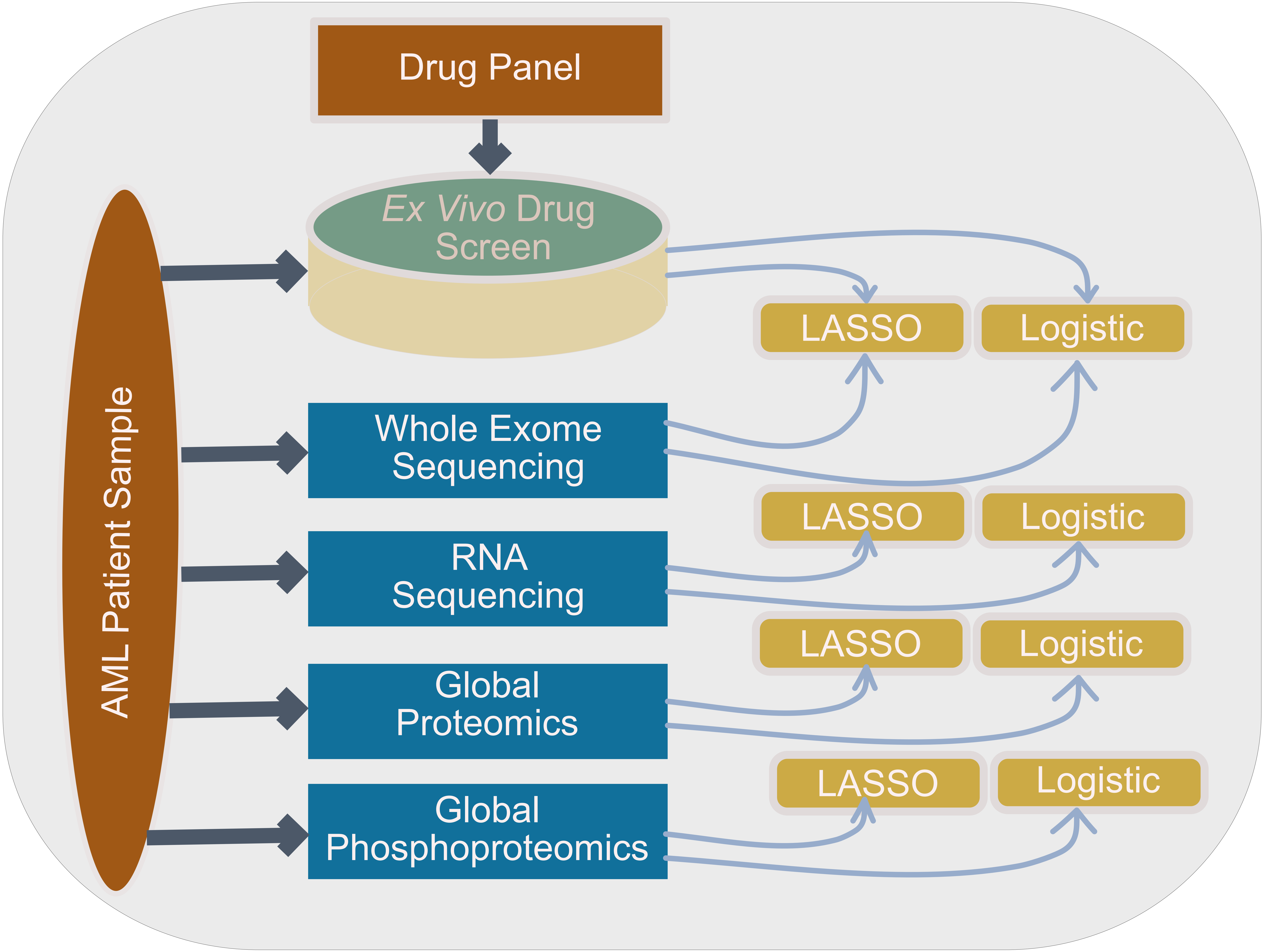

### Supplemental Figure 2

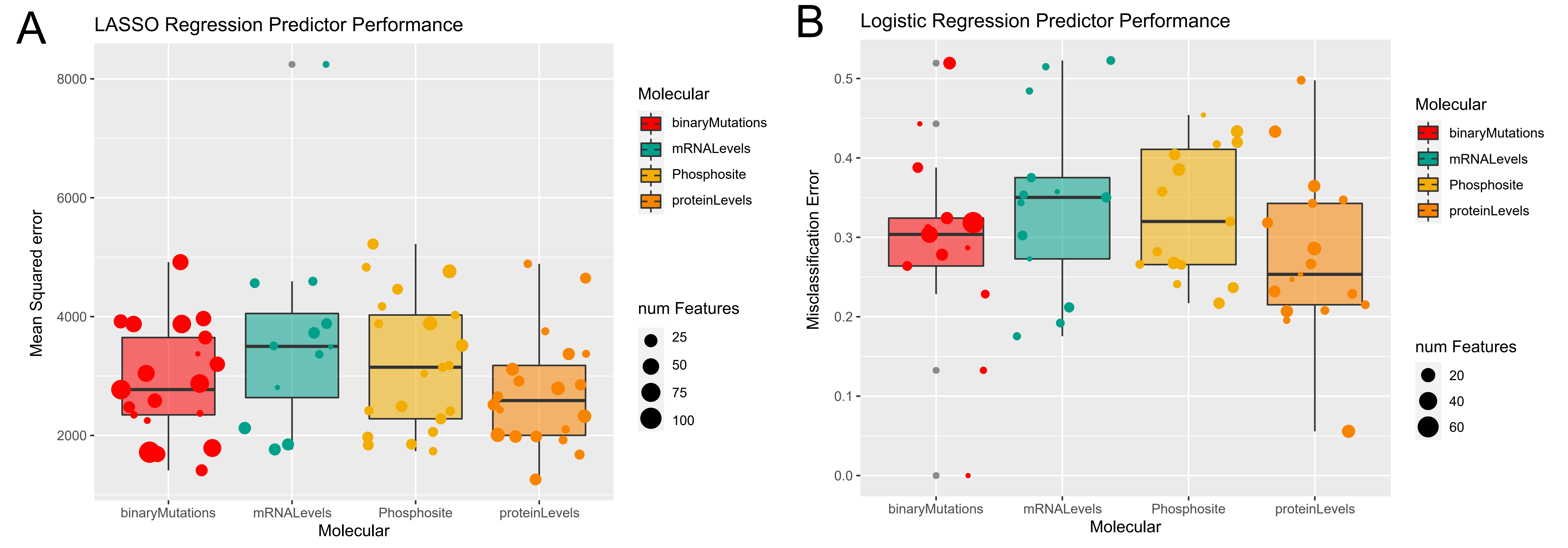

### Supplemental Figure 3

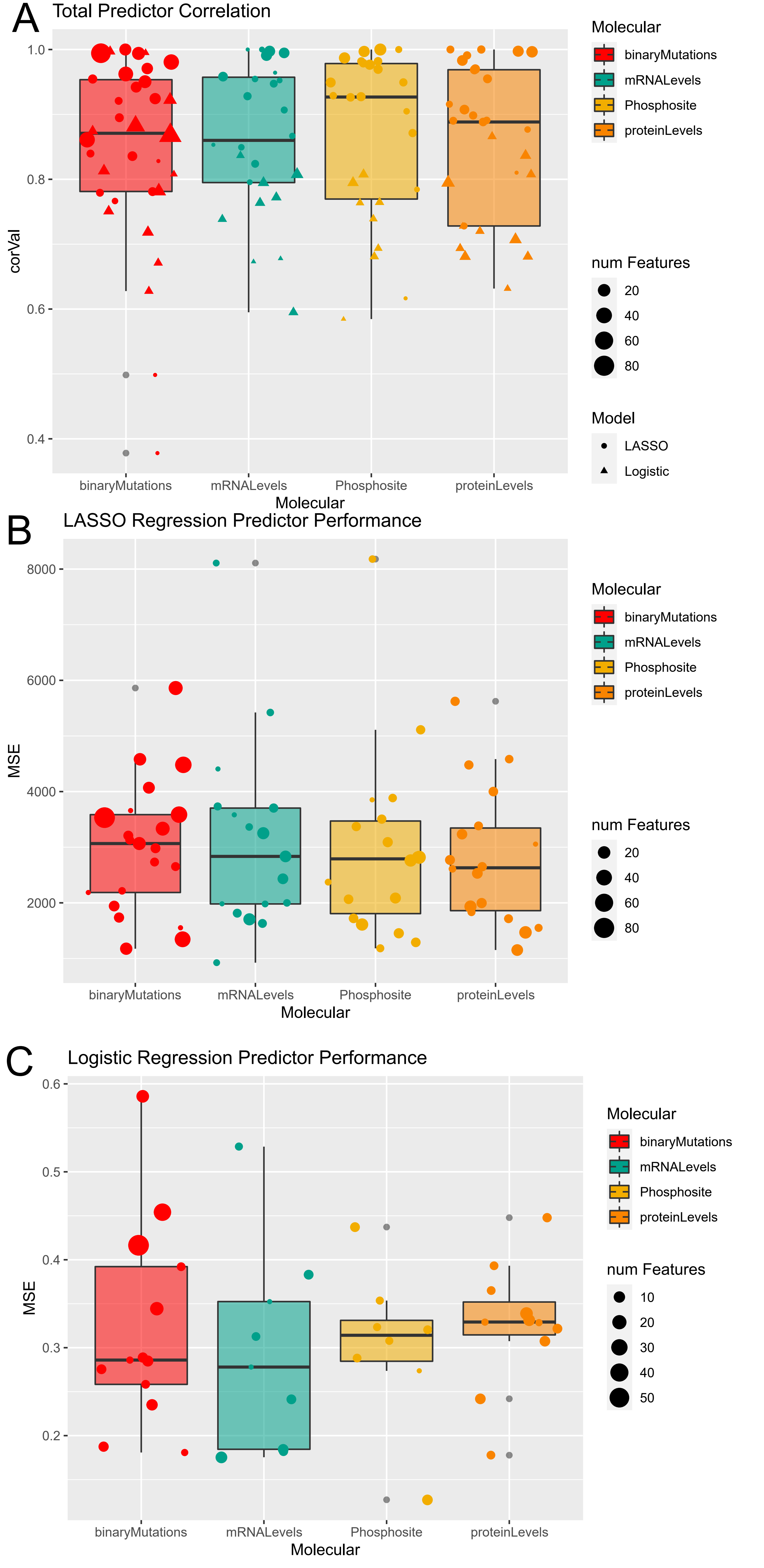

### Supplemental Figure 4

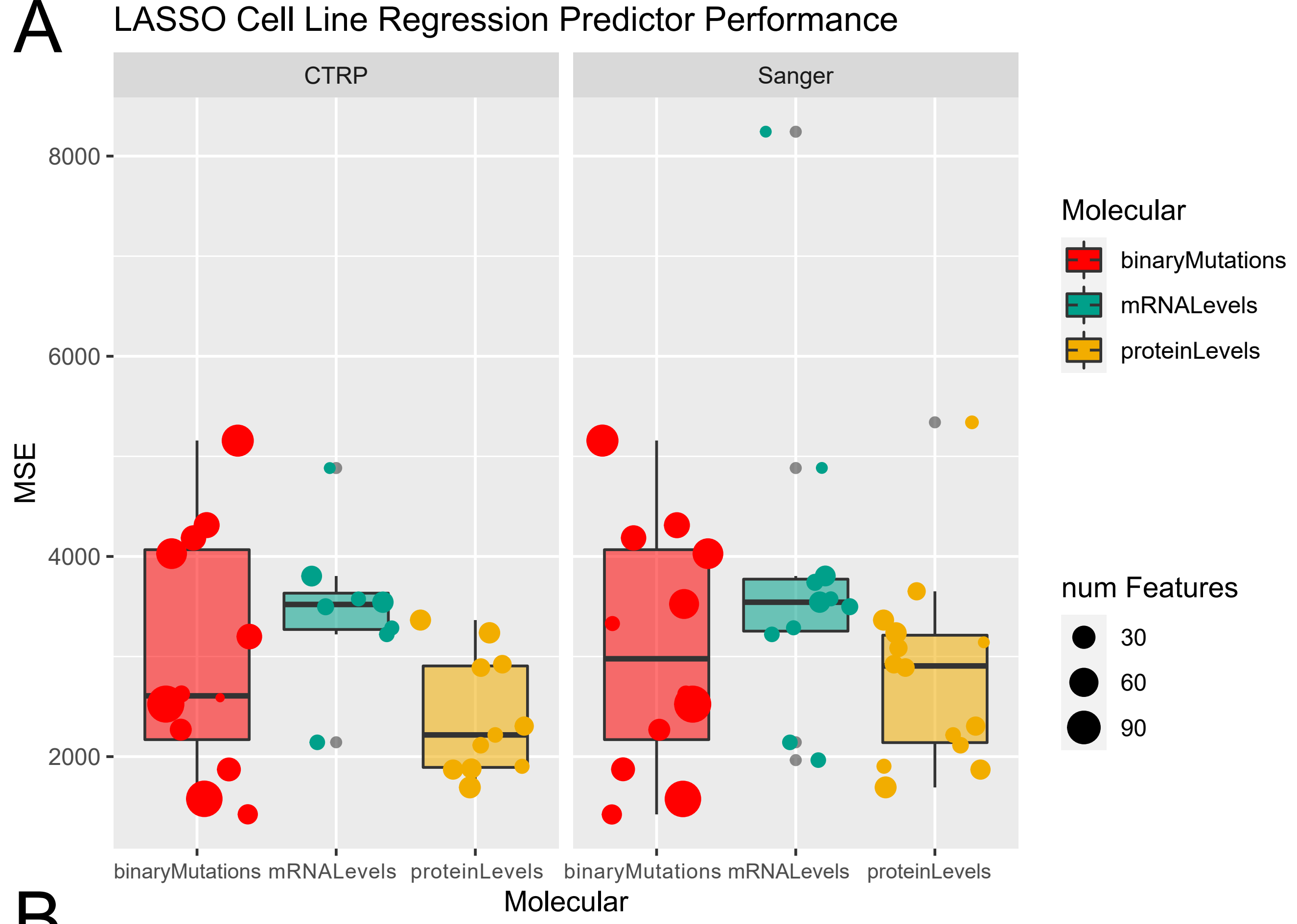

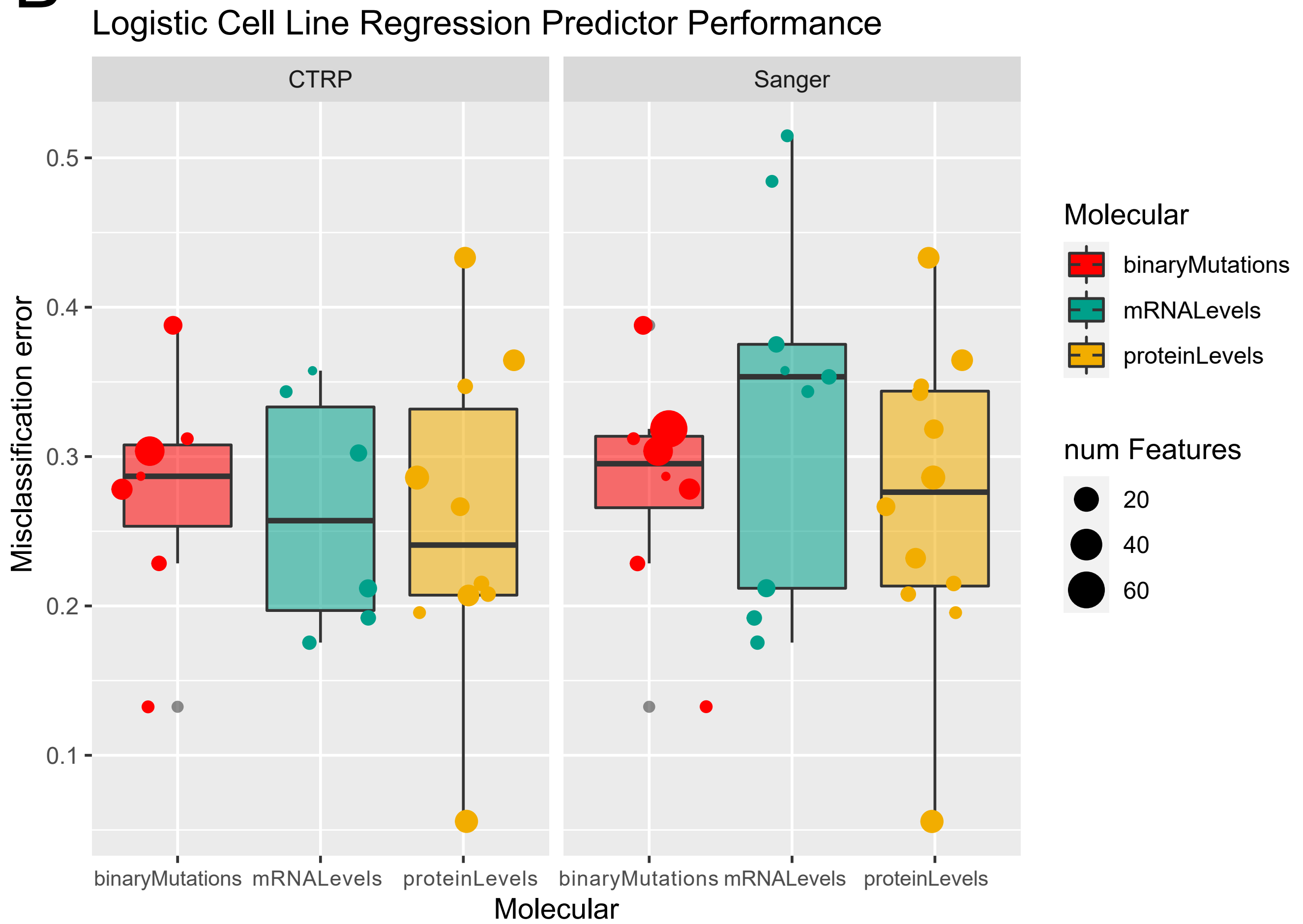
